# Phase Behavior of TDP-43 and hnRNPH1: From Soluble and Aggregated States to Liquid-Like Droplets

**DOI:** 10.1101/2025.11.12.687994

**Authors:** Travis R Wang, Jiwu Wang

**Affiliations:** Allele Biotechnology and Pharmaceuticals, Inc., 6404 Nancy Ridge Drive, San Diego, CA 92121 The Scintillon Institute, 6868 Nancy Ridge Drive, San Diego, CA 92121

**Keywords:** TDP-43, hnRNPH1, Phase Separation, Intrinsically Disordered Region, low-complexity domain, SpyTag-SpyCatcher

## Abstract

Cytotoxic inclusions of TAR DNA-binding protein 43 (TDP-43) have been identified in various neurodegenerative diseases. Mediated by its low-complexity C-terminal domain (CTD), TDP-43 can undergo liquid-liquid phase separation (LLPS) and form aggregates and fibrils. These phase transitions are associated with pathological TDP-43 mislocalization and cytoplasmic aggregation. What determines whether and when TDP-43 adopts a liquid droplet state or forms solid aggregates, transiently or irreversibly? A full understanding of the underlying biophysical mechanisms will help explain the cellular toxicity of aberrant forms of TDP-43 and similar proteins, such as some of the large family of RNA-binding proteins. In this study, we generated recombinant full-length TDP-43 and maintained soluble-protein working solutions without the need of a solubility tag. By focusing on conditions that tune the electrostatic characteristics, we induced phase transitions among states of soluble TDP-43, solid aggregates, and liquid droplet phases in an expedient and controlled manner. We also obtained soluble working solutions of TDP-CTD and another crucial splicing factor, hnRNPH1, and investigated their phase behaviors as individual proteins and their interactions between each other during phase transitions. Using this system, we discovered a previously unreported aggregate-to-droplet transition and found evidence that phase separation of low-complexity domain (LCD)-containing proteins likely involves domain matching when they coaggregate out of solution.

## Introduction

Protein aggregation is a defining feature of many neurodegenerative disorders, including the accumulation of amyloid-β and Tau in Alzheimer’s disease, α-synuclein in Parkinson’s disease, and TDP-43 in amyotrophic lateral sclerosis (ALS). A central question in ALS study is what stimuli trigger TDP-43 aggregation and which elements and properties of TDP-43 respond to the triggering stimuli. TDP-43 is a ubiquitous RNA-binding protein (RBP) that regulates RNA splicing^1^, with roles in regulatory control on mRNA homeostasis, translation, transcription, and transport^2^. Though predominantly residing in the nucleus with an N-terminal nuclear localization signal, TDP-43 shuttles between the nucleus and cytoplasm, a process found to be influenced by cellular conditions such as stress^3,4^. The N-terminus region promotes self-association into dimers and ordered oligomers^5^, with evidence showing oligomerization into higher-order structures prerequisite to elements of protein function^6,7^. Congruent to its role as an RBP, TDP-43 contains two conserved RNA recognition motifs (RRMs) which mediate binding to RNA and DNA and are central for TDP-43’s biological roles. The C-terminus includes a low-complexity domain (LCD) which is glycine-rich and aggregation-prone that regulates protein-protein interactions^8^, splicing, and solubility during physiological function^4,7,9^.

Pathologically, the accumulation of TDP-43 aggregates is observed in 90% of cases of ALS, 50% of frontotemporal lobar degeneration cases, and up to 75% in severe cases of Alzheimer’s disease, according to some estimates^10^. However, even in physiological functions, TDP-43 does not always exist as a free soluble protein; rather, it transitions between solution and reversible liquid-like droplets (LLDs) with limited mobility^3^. These droplets can change in size and shape in response to cellular conditions. It appears that individual TDP-43 molecules and the oligomer complexes they form exist in a constant state of flux; protein-protein interaction and aggregate formation are always in play without necessarily leading to a pathological endpoint.

Under stress conditions, TDP-43 relocates to the cytoplasm to scaffold with other RBPs and RNA into dynamic stress granules, which lock down translation and sequester specific molecules as a coordinated cellular response to stress^11,12^. Extensive studies into stress granule composition have illustrated that increased concentrations of TDP-43 lead to liquid phase separation (LLPS; demixing) and depletion of native protein^13^. When forming different forms of aggregates, the protein concentration and interaction strength among protein molecules are key factors that modulate LLPS from a proteome-wide perspective^14^. One proposed pathway for how soluble TDP-43 congregates and misfolds into toxic aggregates is through prolonged or repeated homotypic-interactions at elevated concentrations within these granules, with other triggers playing a role such as oxidative stress, amino acid mutation and modification, and the presence of RNA^15^. However, studies using tracking and localization microscopy uncovered not only distinct substructures of TDP-43 within stress granules, but also numerous regions of slow-moving TDP-43 species dispersed throughout the cytoplasm, indicating TDP-43 oligomerization and aggregation can also occur outside of stress granules^16,17^, where protein concentrations are not constrained by granule confinement. The precise mechanisms governing the dynamic balance among soluble proteins, oligomer formation, and subsequent transitions into droplets and fibrillar assemblies are subjects of intense study, yet remain poorly understood even with significant progress made over a decade by many groups of researchers.

The term “aggregate” relevant to low-complexity domain (LCD)-containing RNA-binding proteins commonly include oligomers that may be in the form of liquid-like droplets, hydrogel-like “glassy” condensates, or solid formations of various shapes, often progressing in that order ^16–18^. Stress granules and similar membraneless organelles, in which most of this process has been studied, dynamically organize protein–protein and protein–RNA interactions in the cytoplasm and through LLPS segregate into a concentrated phase from a dilute phase. LLPS in proteins is often driven by LCDs, which mediate multivalent interactions. Both full-length TDP-43 and its C-terminal domain (CTD) are known to undergo phase separation in vitro^14,15^.

Insoluble inclusions of TDP-43 proteins containing fragments of its C-terminus are a hallmark in numerous neurodegenerative diseases. It is quite remarkable that nearly all pathology-relevant mutations of TDP-43 reside in its CTD^8^, indicating the sensitivity of CTD’s intrinsic functions to any sequence perturbation despite its disordered structural morphology. This may suggest that, prior to aggregation, factors beyond structural elements, such as the charges of key amino acids, which can be influenced by extensive phosphorylation, acetylation, and other posttranslational modifications concentrated within the region^8,19^. Additionally, charge distribution along the sequence and neighboring effects, including stacking and hydrophobic interactions with specific charged amino acids, may be essential for their behavior during phase transition^20^. In one report, extensive mutagenesis was conducted to identify amino acid sequence as the “molecular grammar” that dictates phase behavior in the Fused in Sarcoma (FUS) family^21^. In another study, adjusting the placement of charged amino acids within an otherwise hydrophobic sequence resulted in a twofold change in the tendency to phase-separate, without altering the net charge of the polypeptide^22^. To manipulate the phase behaviors of TDP-43 through electrostatic alteration, which shares many similarities with FUS, we created conditions that strongly altered the surface charge of the protein and observed that it drove phase transitions into liquid droplets either directly from soluble protein or, unexpectedly, through a novel reversal pathway from aggregates.

Phase separation is typically a thermodynamic process that occurs when protein-protein interactions are more favourable than protein-solution interactions, separating proteins and solutes into a dense phase^23^. Observed TDP-43-rich droplets in cells often contain internal vacuoles filled with nucleoplasm materials, and these coexisting yet immiscible multiphase occurrences correlate with ALS-causative mutations in CTD, indicating a higher-level complexity of phase separation exists and is pathologically relevant^24^. Intriguingly, a reversible double-emulsion structure (droplets-in-droplets) was created in vitro using Poly-rA RNA and PEG and found to be due to kinetic trapping as opposed to thermodynamic equilibrium^25^, but multi-phase droplets have yet to be recreated with TDP-43 or other similar RNA-binding proteins in vitro in a similarly controlled manner. In this report, we present our system as being able to show multi-emulsion droplet formation with simple triggers and a consistent time course.

The CTD of TDP-43 has been found to form LLDs^26,27^. NMR and in vitro assays identified residue 321–335 of TDP-43 in the CTD region to be part of a helical core that tunes LLPS^14^ whereas proteolytic and cryo-EM studies identified residue 279-360 as the fibrillar core of TDP-43 filaments^28^, encompassing the LLPS core. As for full-length TDP-43, phase separation has been reported under unique poly(ADP-ribose) (PAR) binding conditions with concentrations of TDP-43 at 10µM^29^, though in vitro LLPS of the full-length protein is still difficult to induce or control even with favorable conditions such as using crowding agent PEG or aggregation-inhibiting molecules dopamine and polyphenol EGCG^30,31^. Here, we present a novel system of manipulating liquid phase separation of CTD and the full-length TDP-43, and for comparison, hnRNPH1. The current paradigm states that TDP-43 LLPS droplets age into hydrogels, inclusions, and amyloid fibrils. LLPS acts as a scaffold accelerating solid aggregate formation, often leading to direct amyloidosis^30^. Our hypothesis is that the phasing-out of the soluble form starts with a “seed” area in a protein-specific manner, which leads to early-stage substructural fibrils, forming solid aggregates or LLDs depending on the predicted charge of this region and subsequent electrostatic interactions. We further show the protein-specific seeding results by using switchable combinations of fluorescent labelling schemes.

Our findings represent an unusual reversal of the conventional uni-directional liquid-to-solid maturation of protein condensates, showing a more dynamic nature of the three macroscopic phases of LCD-containing RNA-binding proteins, and reveal the likely role of electrostatic charge in regulating LLPS.

## Results and Discussion

### Multidirectional phase transitions of TDP-43 are reversible and tunable

Liquid–liquid phase separation of full-length TDP-43 has been viewed as difficult to recreate in vitro using recombinant proteins^32^. Several reported cases relied on an enzymatic cleavage of a solubility tag to initiate aggregation, which is typically monitored by increased turbidity with the enzymatic reaction difficult to control^7,33–35^. In other systems, a solution jump triggers phase separation either almost instantly or very slowly—within minutes after a solution jump^29^, or over 5 days^28^. In all cases, it has been challenging to fine-tune the process and capture time points of phase transitions. To better control the transition process to gain clearer insight into the phase behaviors of these proteins, we sought to generate recombinant full-length TDP-43 of sufficiently high soluble concentration and stability without a large solubility tag.

The first step was to choose a method to keep TDP-43 soluble through purification. Most previous attempts to produce full-length TDP-43 have relied on large solubility tags, such as GST^36^, MBP^37^, and SUMO^15,28,31^, which keep the fusion protein soluble until aggregation is triggered by enzymatic tag cleavage or solution jump. Alternatively, like all proteins, recombinant TDP-43 can be readily solubilized using strong denaturants such as 6-8 M urea or guanidine, with or without a solubilizing tag, and this method has been employed by different investigators^29,36^. However, as complete removal of these denaturants is often problematic because the protein tends to precipitate upon denaturant elimination, a residual low concentration (e.g., ∼0.6 M urea or guanidine) is typically retained in storage buffer after purification, which may interfere with subtle aspects of phase behavior. For our in vitro phase analysis system, we aimed to establish a baseline using full-length TDP-43 without interference from large solubility tags or residual denaturants. We chose to produce full-length TDP-43 with only a small His6-tag for purification and use urea to keep it soluble during production, then refold to urea-free storage buffer. We solubilized full-length TDP-43 expressed in E. coli using 6M urea that completely denatured its secondary structures without affecting affinity purification via the His-tag. After washing and eluting with imidazole from cobalt beads in 6 M urea-containing buffers, urea was gradually removed by stepwise dialysis against buffer at 100-fold the sample volume. The urea concentration was reduced by half every 1–2 hours over 5-6 steps, concluding with a final exchange into urea-free buffer.

As the second step, we formulated the protein storage buffer by first testing whether high or low salt conditions promote solubility. In the field of membraneless organelles and coacervates, critical salt concentration is determined by the intrinsic properties such as charge-to-neutral ratio among monomers of polymeric molecules such as DNA, RNA or protein, and charge distribution and pattern^22,38^. Reducing the salt concentration below the critical concentration leads to energy-favorable segregation of many types of proteins. This would suggest that keeping salt concentration high should avoid forming aggregates during purification steps and in storage. Many previous studies indeed used storage buffers containing high salt, e.g. 0.5 M NaCl or KCl^28,31,36,37^. However, we found that TDP-43 tends to aggregate more under high-salt conditions (e.g., 0.5 M NaCl) than under low-salt conditions (Supplementary Fig. 1a). Recent studies have also reported that high salt promotes phase separation of soluble TDP-43^29,37^.

The observation that high salt destabilizes the soluble state of TDP-43 suggests that charge screening enhances hydrophobic and other non-ionic interactions, thereby promoting phase separation. We hypothesized that similar effects could be achieved by adjusting the buffer pH toward the protein’s isoelectric point (pI). To test this, we prepared TDP-43 (5 μM) in a low-capacity pH buffer (20 mM Tris-HCl, pH 8.0) and low salt (100 mM NaCl) and titrated the pH by adding 1 M Tris-HCl at the target pH, maintaining 100 mM NaCl. Approximately 10% of the final volume was required to reach the desired pH. Under these conditions, TDP-43 remained soluble at pH 8.0, but the solution became turbid with diffuse assemblies at pH 7.0. At pH 6.0—near the pI of full-length TDP-43—distinct solid aggregates became evident (Supplementary Fig.1b). This observation is consistent with findings from the Chiti group^29^, which implied that electrostatic repulsion normally inhibits phase separation. Based on these results, we set the storage buffer for recombinant TDP-43 as 20 mM Tris-HCl, pH 8.0, 100 mM NaCl.

By including urea during purification with a solubility-friendly storage buffer, we obtained working solutions of TDP-43 suitable for phase transition analysis. A few points are noteworthy: a stepwise removal of urea in our protocol enables the protein to refold through a more controlled process, rather than by dropwise dilution or leaving residual denaturant^29,36^. Complete denaturation with urea also eliminates the need for extensive RNA digestion followed by ion-exchange and gel-filtration chromatography to remove nucleotides, which, if retained, can interfere with phase separation^35^.

To initiate analysis of the solution-to-aggregation transition, we altered salt concentration and pH by adding 5 M NaCl and 1 M Tris-HCl (pH 6.0) to aliquots of TDP-43 (5 µM) in storage solution. Once the buffer composition changes to high salt (0.5 M NaCl) at pH 6.0, TDP-43 almost immediately aggregated out of solution and formed solid globules (Fig 1A). Over several hours, these globules gradually assembled into larger structures composed of subunits that merged into similarly disordered arrangements, reminiscent of fractal-like architectures (Fig. 1a). After 3 hours, these aggregates ranged from clusters of only a few smaller congregates to assemblies spanning hundreds of microns. The appearance of these structures closely resembles those described by Lashuel and colleagues, which were composed of full-length TDP-43 oligomers released from immobilized metal affinity chromatography only in high-imidazole (500 mM) fractions^28^. Unlike our process, their assemblies required approximately five days to form. To assess whether a commonly used solubility tag would alter TDP-43 phase behavior, we tested SUMO, a 101-residue natural fragment with a pI of 5.2. When fused to TDP-43, the resulting construct has a pI of 5.6, close to untagged TDP-43, leading us to predict that both versions would exhibit comparable phase properties under our pH- and salt-based phase transition conditions. We prepared a SUMO–TDP-43 fusion using the same 6 M urea denaturation and renaturation protocol and observed very similar phase behavior to untagged TDP-43 (Supplementary Fig. 2a). These results suggest that SUMO tagging preserves TDP-43 phase characteristics and may be suitable to study phase transitions of other proteins, although pI matching should be considered.

**Figure 1.**
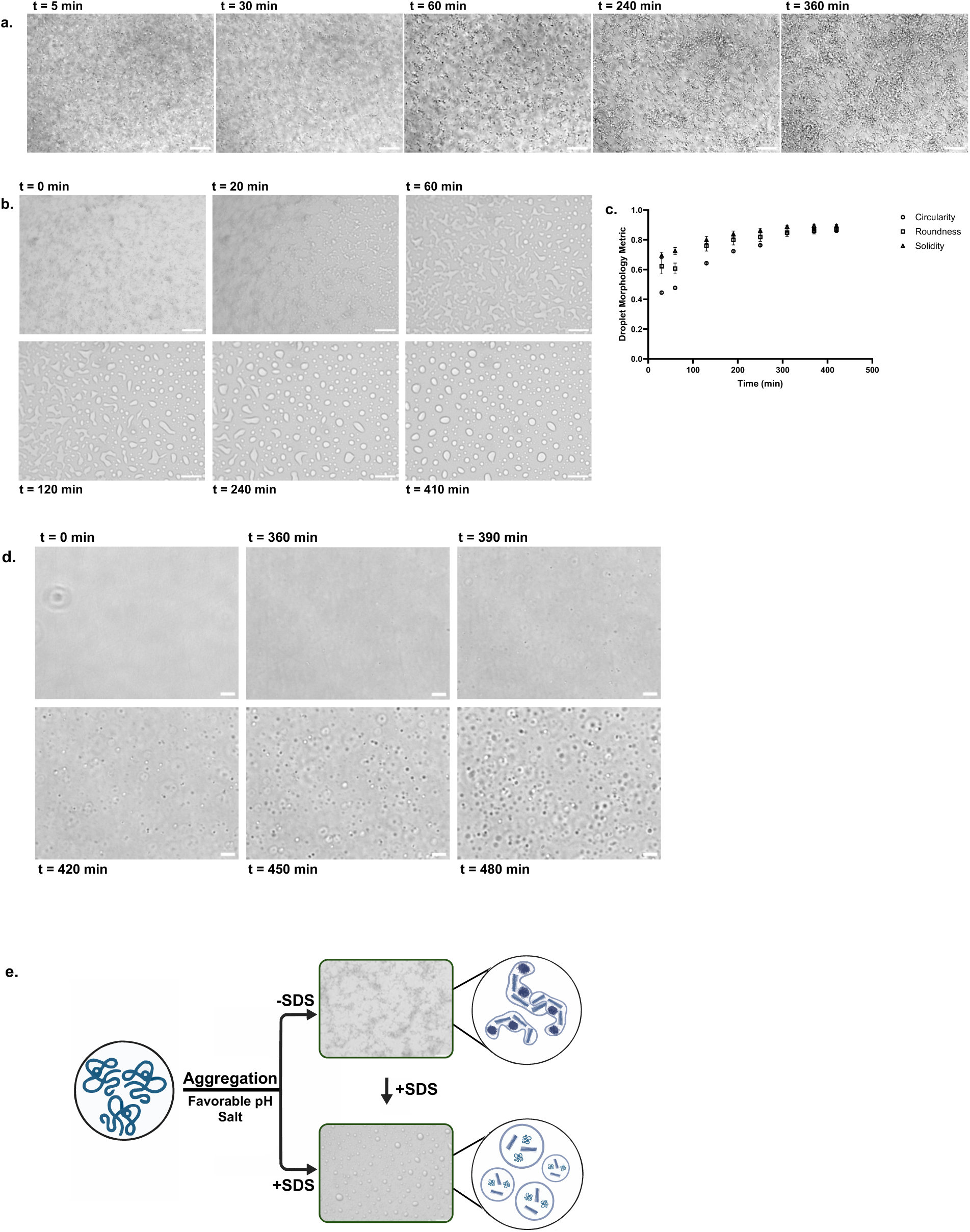
Three-phase transition of recombinant full-length TDP-43 induced by charge modulation. **a** Gradual formation of solid aggregates from soluble TDP-43 at pH 6 and 0.5 M NaCl. Scale bar is 40μm. **b** Solid aggregates converted into a hydrogel upon addition of SDS (final concentration 0.03%), then progressively transformed into spherical droplets over 7 hours. Time notations begin at the addition of SDS. Scale bar is 40μm. **c** The diagram illustrates the time-dependent morphological changes measured by circularity, roundness, and solidity (see Materials and Methods). Error bars are Standard Errors of the Mean. **d** Droplets formed directly from solution following SDS addition, with the most pronounced transition occurring between the 6th and 8th hour. Scale bar is 20μm. **e** Schematic representation showing how TDP-43 can transition among all three phases through controlled induction.

Our findings that a change in TDP-43 net charge can modulate aggregate formation prompted us to investigate whether alternative strategies for manipulating charge could similarly influence TDP-43 phase behavior. The strong anionic detergent sodium dodecyl sulfate (SDS) micelles facilitate denaturation of proteins and destabilization of protein complexes. The extent of destabilization depends on the SDS concentration and the protein−SDS stoichiometry and the resistance to 0.1-2% SDS has been used as a test of TDP-43 fibrillation strength^31,39^. Because SDS introduces negative charges along the entire polypeptide, we therefore employed SDS at a very low concentration, 0.03%, as a means of mildly modulating the overall protein surface charge. Upon addition to TDP-43 aggregates, strikingly, the solid aggregates dispersed in the field of view began to ‘melt’ into glassy material, but instead of dissolving and disappearing, progressively transforming in shape and eventually forming round condensates, in about 6–7 hours (Fig 1b). Formed droplets were quantified in circularity, roundness and solidity using ImageJ quantification functions (Fig 1c). Coating aggregated TDP-43 with negative charge via SDS did not dissolve the aggregates into a soluble state but instead transformed them into droplets. We next asked whether introducing net negative charge to soluble TDP-43 would directly induce LLPS. As shown in Fig. 1d, adding SDS to soluble TDP-43 indeed triggered droplet formation, although the droplets were smaller and less dense than liquid-like formations formed from aggregates. These early-phase droplets could be further enlarged and increased in abundance by adding PEG8000 (Supplementary Fig. 3), a crowding agent commonly used to promote LLPS^29,36^. The transitions from soluble state to solid aggregates induced by pH and salt, and from soluble state or, as a novel observation, from solid aggregates to liquid-like droplets upon SDS addition are schematically illustrated in Fig 1e.

The current paradigm for phase separation of LCD-containing RNA-binding proteins describes a progression from soluble molecules to liquid-like condensates, then to glassy solids, and ultimately to amyloid fibers^18^. It is interesting to compare the SDS-induced LLDs with LLDs formed by means of adding negative net charges other than SDS. For instance, phosphomimetic mutations reduced TDP-43 aggregates and increased the appearance of larger, more liquid-like droplets^33^. Inside cells, phosphorylation is expected to add negative charges and there are many other factors that can alter the net charge and charge distribution of proteins. McGurk *et al.* reported that non-covalent binding of TDP-43 to poly(ADP-ribose) (PAR), a negatively charged biopolymer and cellular regulator of protein localization and liquid demixing, promotes TDP-43 LLPS in vitro and is essential for its accumulation in stress granules in neurons^15^. Non-covalent binding to PAR resulted in droplets comparable to those in our system using SDS, likely from similar effects of adding net negative charge to TDP-43. As a matter of fact, binding to RNA would add net negative charge to TDP-43 as well in normal cellular environment, which could enhance LLPS according to our model along with suggestions by others^40^. Although SDS has not, to our knowledge, been used as a trigger to induce LLPS in TDP-43 or other LCD-containing RNA-binding proteins, it has been reported to promote LLPS of amyloid-β oligomers at supraphysiological concentrations^41^, suggesting its broader potential as a tool for modulating phase behavior across diverse proteins. Demonstrating reversibility of solid formations with strong anionic modulators could shed light on understanding TDP-43 pathology and lead to the development of new therapeutic strategies.

### SDS-induced droplets possess typical liquid-like properties

Three lines of evidence indicated that droplets derived from solid aggregates shown above exhibit the same liquid properties as those formed by LLPS. First, these droplets fuse upon close contact (Fig. 2a). Second, they dissolve in 1,6-hexanediol, similar to typical LLPS droplets (Fig. 2b). Recent work has shown that glycine and other amino acids can bind phase-separating proteins and modulate condensate stability and liquid-like dynamic^42^, therefore, as a third test, we added glycine to the droplets at 50 mM and observed pronounced changes within minutes: namely, a dynamic formation of droplets-in-droplets, subsequent growth, and eventual remerging over time (Fig. 2c). These dynamic changes were quantified by tracking the number of internal droplets relative to the size of the encompassing outer droplets (Fig. 2d), as well as the average count of droplets over a 12-minute period (Fig. 2e). Upon glycine addition, there was a rapid increase in internal droplet formation, coordinated with the growth of the encompassing droplets. This was followed by a consolidation phase, during which the average number of internal droplets decreased. Altogether these corroborating assays provide strong evidence that the dynamic liquid-like behavior of these solid-derived droplets.

**Figure 2.**
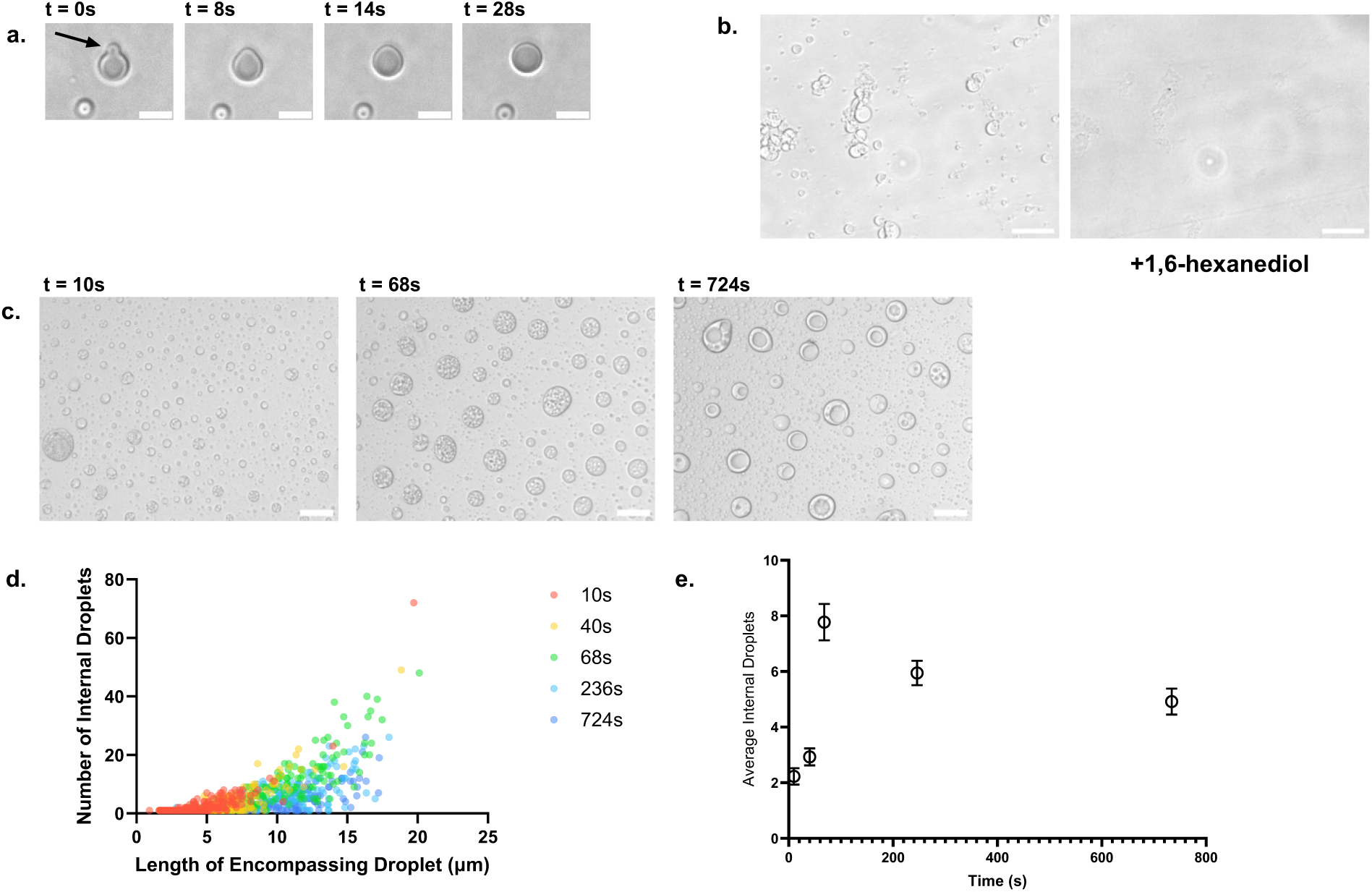
Droplets induced from solid aggregates exhibit liquid-like properties. **a** Time-lapse images showing a small droplet (arrow) gradually merging with a larger droplet. Scale bar is 10µm. **b** Droplets derived from aggregates dissolve upon treatment with 1,6-hexanediol, consistent with typical LLPS behavior. Scale bar is 40μm. **c** Formation of multi-emulsion droplets upon addition of glycine (50 mM) to initial aggregate-derived droplets. Time indications begin at addition of glycine. Scale bar is 20μm. The transition occurs on the scale of minutes, progressing from small droplets (left) to multiple inner droplets encapsulated within larger outer droplets (middle), and finally to fused, larger droplets (right). **d** Graph showing the number of internal droplets (Y-axis) versus the size of the encompassing droplet (X-axis) and time in seconds (color legend). **e** Graph displaying changes in average number of internal droplets per encompassing droplet (Y-axis) over time in seconds (X-axis). Error bars are Standard Errors of the Mean.

When Cy5-labeled mRNA of a fluorescent protein mWasabi as biologically irrelevant mRNA was allowed to bind TDP-43 prior to phase separation, the Cy5 imaging revealed that the mRNA colocalized with the protein within the dense phase of the multi-emulsion droplets (Supplementary Fig. 4), consistent with previous findings that full-length TDP-43 oligomers retain RNA-binding capability^31^. These observations further indicate that the contents of these droplets exhibit liquid-like properties similar to typical LLPS condensates, despite originating from solid assemblies transformed by forced charge alteration.

### Phase transition behavior of TDP-43 can be generalized to other RNA-binding proteins

To see if any of these phase behaviors are specific to full-length TDP-43, we wanted to compare TDP-43 to hnRNPH1, another nuclear RNA-binding protein that regulates alternative splicing with implications in ALS and other diseases^43^. Similar to TDP-43, hnRNPH1 also has an LCD at the C-terminus of the polypeptide composed of 449 residues, close to 414 in TDP-43; its pI is the same at pH 6, allowing us to apply the same pH and salt conditions to manipulate the electrostatic and ionic environment for phase transitions as TDP-43. We also wanted to compare the whole TDP-43 protein to its aggregation-prone domain. In vivo, neuronal inclusions contain ubiquitinated and phosphorylated aggregates of TDP-43 C-terminal (TDP-CTD) fragments in various sizes (e.g., TDP-25 and TDP-35), created by aberrant cleavage events that contribute to ALS^44,45^. TDP-CTD was initially identified as the fragment encoding amino acids 274-414 by pioneering studies^14,26^. Later, 266-414^46^, 267-414^27^, and longer CTD 193-414 were also studied^47^, and all CTD segments have been found to play a major role in phase separation as well as aggregations of TDP-43. The 274-414 fragment CTD has a pI close to pH 10. It is previously reported to have a different phase separation response to pH gradients from the full-length^29^, likely related to different net charge at different pH compared to the full-length TDP-43 protein. We therefore opted to study an intermediate-sized version of TDP-CTD, spanning 208 to 414 amino acids, which has a predicted pI at pH 6. This allowed us to compare it to the full-length TDP-43 in parallel and during protein mixing experiments.

As with full-length TDP-43, both TDP-CTD (208-414, hereby referred to as TDP-CTD in this report) and hnRNPH1 formed solid assemblies when their solutions were adjusted to pH 6.0 and 0.5 M NaCl. Upon addition of 0.03% SDS, these assemblies transitioned into droplets, exhibiting behavior broadly similar to that of full-length TDP-43 (Fig. 3a). However, the dynamics and morphological details of the resulting aggregates and droplets varied among the proteins. For example, TDP-CTD consistently formed larger and more compact aggregates compared to TDP-43 and hnRNPH1. Whilst the primary focus of this report is to demonstrate macroscale phase transition processes that can be modulated by solution conditions, further insights into the fine structural differences underlying the various assembly forms of these proteins could be gained by high-resolution imaging in future studies. This result nevertheless supports that the pI and high salt promotion of solid aggregation and their SDS-induced conversion into droplets are not specific to full-length TDP-43. This is consistent with the notion that phase behavior can be dictated by a general mechanism based on electrostatic and hydrophobic interactions.

**Figure 3.**
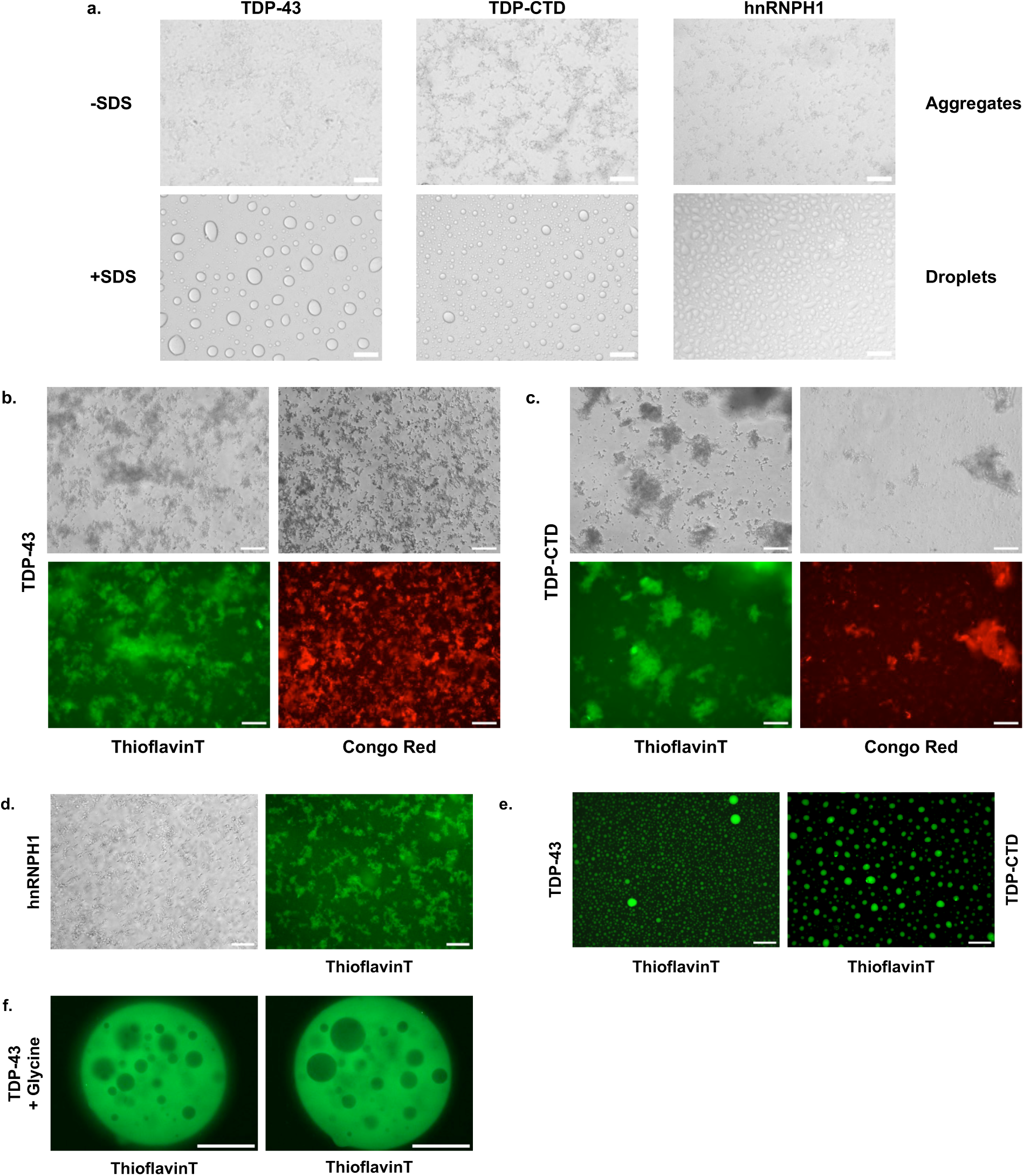
TDP-43, TDP-CTD, and hnRNPH1 form amyloid-positive aggregates and droplets. **a** Side-by-side comparison of solid aggregates formed by inducing soluble TDP-43, TDP-CTD, and hnRNPH1 at pH 6.0 and 0.5 M NaCl (top), and droplets formed after further addition of SDS to a final concentration of 0.03% (bottom). Images were taken 8 hours after induction. Scale bar is 20μm. **b-d** Aggregates formed by TDP-43 (**b**) and TDP-CTD (**c**) stain positive with ThT and Congo Red (top: bright field; bottom: fluorescence images). **d** hnRNPH1 also stains positive for ThT. Scale bar is 40μm. **e** Representative data showing droplets formed directly from soluble TDP-43 and TDP-CTD solutions exhibit ThT staining. Scale bar is 40μm. **f** Multi-emulsion droplet formed upon addition of glycine (50 mM), with ThT included; shown are confocal fluorescence images from a representative Z-stack acquired using Cytation10 (40×). The z-distance between the two slices is 20μm. Scale bar is 100μm.

TDP-CTD droplets and aggregates are known to contain amyloid filaments^27,48,49^. To examine the amyloid nature of our TDP-43 and TDP-CTD solid aggregates, we stained them with classical amyloid-binding dyes, including thioflavin-T (ThT) and Congo Red (CR), both of which strongly fluoresce upon binding to β-sheet-rich amyloid structures. They exhibited clear and nearly uniform positive staining, suggesting that amyloid structures are present throughout the formations and likely organized at the substructural level rather than being confined to larger, assembled structures (Fig 3b). Next, we stained the droplets formed from both TDP-43 and TDP-CTD solid aggregates with SDS addition and discovered that these droplets too were remarkably ThT and CR positive (Fig 3c). Extensive studies by many groups on LLPS of TDP-CTD have shown that amyloid fibrillation can occur independently of LLPS, as observed in our solid aggregates; whereas ThT-positive amyloid fibrils have also been detected within liquid droplets, consistent with our findings in solid-derived droplets^27,50^. Although dye-staining with ThT and CR by itself is not unique to amyloids, the strong and consistent signals here nevertheless provide clear likelihood that these assemblies contain amyloid elements and justify further investigations using high-resolution imaging techniques when available for analysis of the assembly dynamics. Namely, how the helical structures present in both full-length TDP-43 and its CTD during assembly transitions can potentially give rise to β-sheet-rich structures^26,39,46^.

The formation of multi-emulsion or multiphase droplets is a characteristic feature of complex coacervate (polyelectrolyte material) mixtures, which are widely viewed as models for studying charge-mediated LLPS^38^. By patterning charged monomers along a polymer backbone such as polypeptides, one can precisely adjust local charge organization and the resulting electrostatic interactions between coacervate-forming chains, creating multi-phase or multi-emulsion droplets^22^. In TDP-43, charged amino acids are naturally distributed along the polypeptide, with additional charges that can be introduced in vivo through phosphorylation and other post-translational modifications, particularly concentrated in the CTD. In our in vitro phase-separation system, a droplets-in-droplets morphology can be induced simply by adding glycine to SDS-induced droplets (Fig 2c). As previously reported, glycine interacts weakly with amide groups in the protein backbone and aromatic side chains, reducing backbone–backbone contacts while enhancing interactions among aromatic residues^42^. The biological relevance lies in the surprising observation that free amino acids account for more than 25% of the total volume and 6% of the total dry mass of a mammalian cell^51^, as well as their natural role in stabilizing proteins and colloidal dispersions^52^. We set out to determine the nature of multi-emulsion droplets formed following glycine addition to SDS-induced single droplets. To do this, we repeated the experiment shown in Fig. 2c but in the presence of ThT, and used confocal imaging to visualize phase behavior and amyloid properties. Z-stacks of representative complex droplets revealed alternating dense (coacervate) and dilute (supernatant) phases within larger droplets (Fig. 3d). The ‘bubbles’ were ThT-negative, whereas the coacervates were ThT-positive, confirming the presence of amyloid substances in protein-rich coacervates at the 3D level. This result further highlights the liquid-like dynamics of SDS-induced droplets and the secondary phase separation that can be triggered by electrostatic effectors such as glycine.

### Amyloid formation starts upon phase separation

Compared to the CTD of TDP-43, findings reported in the literature on the phase separation behavior of full-length TDP-43 and the role of amyloid fibrillation are less consistent, presumably due to the challenges of producing recombinant full-length TDP-43 without large solubility tags or denaturing agents such as urea, even at low concentrations^32^. Among the few studies on full-length TDP-43, some reported the presence of aggregates composed of proteins with disordered secondary structure that failed to bind ThT^19,53,54^. Other studies observed ThT-positive segregated structures, but the process required extended time, approximately 10 hours in one case^55^ and 9 days with agitation in another^56^. Understanding this time course may provide valuable insights into the fibrillation pathway, including the involvement of LLPS and solid aggregates, and is essential for clarifying how disease-associated amyloid fibrils form. Leveraging the controllability of aggregate and droplet formation using our solubility tag-free recombinant proteins and their strong ThT staining once high order structures form, we performed fluorescence video recording to capture the dynamic process of TDP-43 solutions transitioning into droplets with SDS or forming solid aggregates under conditions of pH near the pI and high salt. Time-lapse images extracted from videos recorded at fixed fields of view from two independent repeats show that, upon addition of SDS to 0.03% at time zero, small droplets began to emerge and progressively increased in size, through joining from soluble proteins or by merging with other condensates (Fig. 4a). Note that ThT was included in the protein solution prior to SDS addition and showed positive staining in both small and large droplets at the first captured time point. While fluorescence intensity increased over time, the rate of increase appeared generally in line with the growth in droplet size, with a slight delay that could reflect amyloid fibrillation progressed in addition to droplet growth; the total droplet counts increased during the first 4 hours before decreasing through merging (Fig. 4b). A low concentration of glycine at 5mM, 10% of what was used in Fig 2c, was also included, and by hour 7, double-emulsion droplets shown as dark circles began to appear, consistent with the results shown in Fig. 2c but at a slower pace. We then applied the same video recording approach to monitor solid aggregate formation from soluble solutions. Similar to SDS-induced droplet formation, aggregates triggered by pH near the isoelectric point and high salt showed ThT staining shortly after the induction. These amorphous, fractal-like solids continued to grow, originating entirely from recruiting soluble proteins as no merging events were observed (Fig. 4c). Analysis of the area and intensity of the solid formations supports our interpretation that aggregates rapidly grew from unstained protein molecules and began to plateau at around 3 hours (Fig. 4d). These observations were consistent across multiple trials and variations using different imaging systems, all showing a smooth transition from unstained solution to clear and immediate amyloid-positive staining. This behavior contrasts with some of the previous reports on full-length TDP-43, as discussed above. The discrepancy may stem from our approach: our TDP-43 protein was fully denatured and then slowly renatured during recombinant protein preparation, likely maintaining a more native-like conformation ready for orderly assembly. Additionally, our induction was carried out by adding the triggering solution at less than 10% of the final volume, creating lower perturbation or stress to the system than other protocols by dropping high concentration protein solutions into low concentration buffers, perhaps better reflecting biological events in vivo.

**Figure 4.**
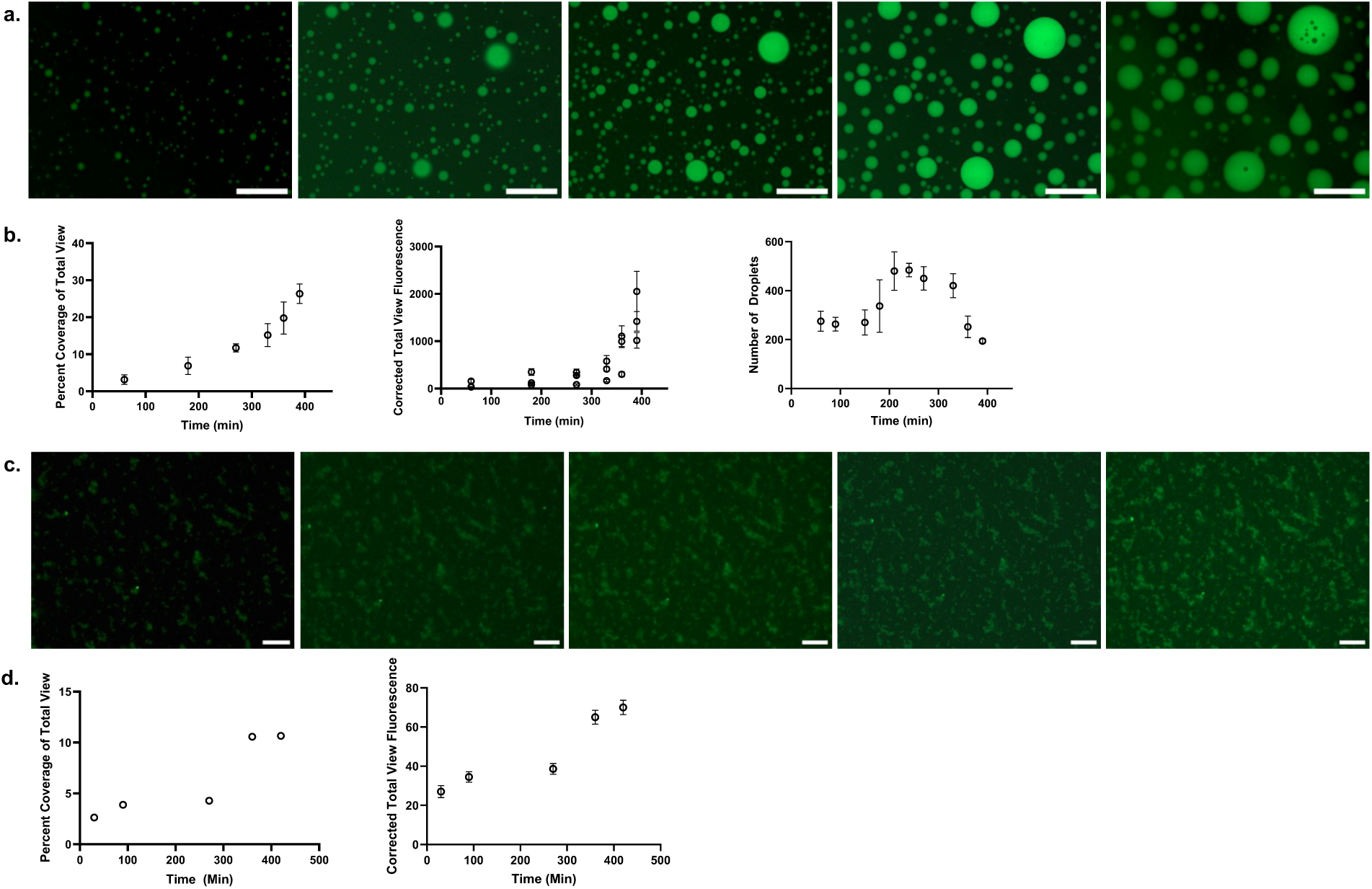
Time course of amyloid-containing droplet and solid aggregate formation. **a** Still photos taken at indicated time points from a time-lapse series of fluorescence image capturing droplets formed from soluble TDP-43. ThT was added to protein solution before addition of SDS (0.03% final concentration) at time 0. Arrows point to example droplets in the merging process. Scale bar is 100μm. **b** Quantitative analysis of count, percentage of view covered, and Corrected Total View Fluorescence (CTVF) of the droplets formed from 3 separate, non-overlapping views of the same time lapse course. Droplet count and area covered are averaged from the 3 views while CTVF is presented from each time point and view. **c** Still photos taken at indicated time points from a time-lapse series of fluorescence image capturing solid aggregates from soluble solution of TDP-43. ThT was added to protein solution before addition of 1M Tris-HCl pH6 and 5M NaCl at 10% final volume at time 0. Scale bars are 50μm. All reactions were carried out at room temperature, ∼25°C. **d** Quantitative analysis of view covered and fluorescence intensity of the solid aggregates formed. Data came from 1 representative view of the time lapse.

### Switchable fluorescent protein tags are a valuable tool for studying protein phase behavior

We were also interested in determining whether different proteins, sharing the same or similar but distinct LCDs, would co-aggregate or remain segregated upon triggering phase separation. To address this, we designed a system to track multiple proteins using distinguishable fluorescent labels. Because phase separation in our system depends on charges that influence electrostatic and hydrophobic interactions, we avoided commonly used dyes that bind to amino acids and may alter charge. Structural dyes such as ThT and CR are unsuitable for aggregate combination studies because they lack protein-specific tracking capability. Instead, we adapted a switchable tagging system to label TDP-43, TDP-CTD, and hnRNPH1 with spectrally distinct fluorescent proteins (FPs). A 13-amino-acid SpyTag was added to the C-terminus of each protein via a 7-amino-acid linker, and the 116-amino-acid SpyCatcher was fused to the N-terminus of mNeonGreen^57^, mScarlet3^58^, and mTurquoise2^59^, all of which represent FPs with optimal brightness, physical and photo stability, and more stringent monomeric properties within their respective spectrum to maximize fluorescence signals while minimizing aggregation artifacts (Fig. 5a). By briefly mixing a pair of SpyTag- and SpyCatcher-fused proteins, a fluorescent label is covalently attached to the target protein via SpyTag–SpyCatcher bonding. The protein ratios within each pair were optimized, with a slight excess of SpyCatcher-FP to ensure that all SpyTags on target proteins were occupied (Supplementary Fig. 5).

**Figure 5.**
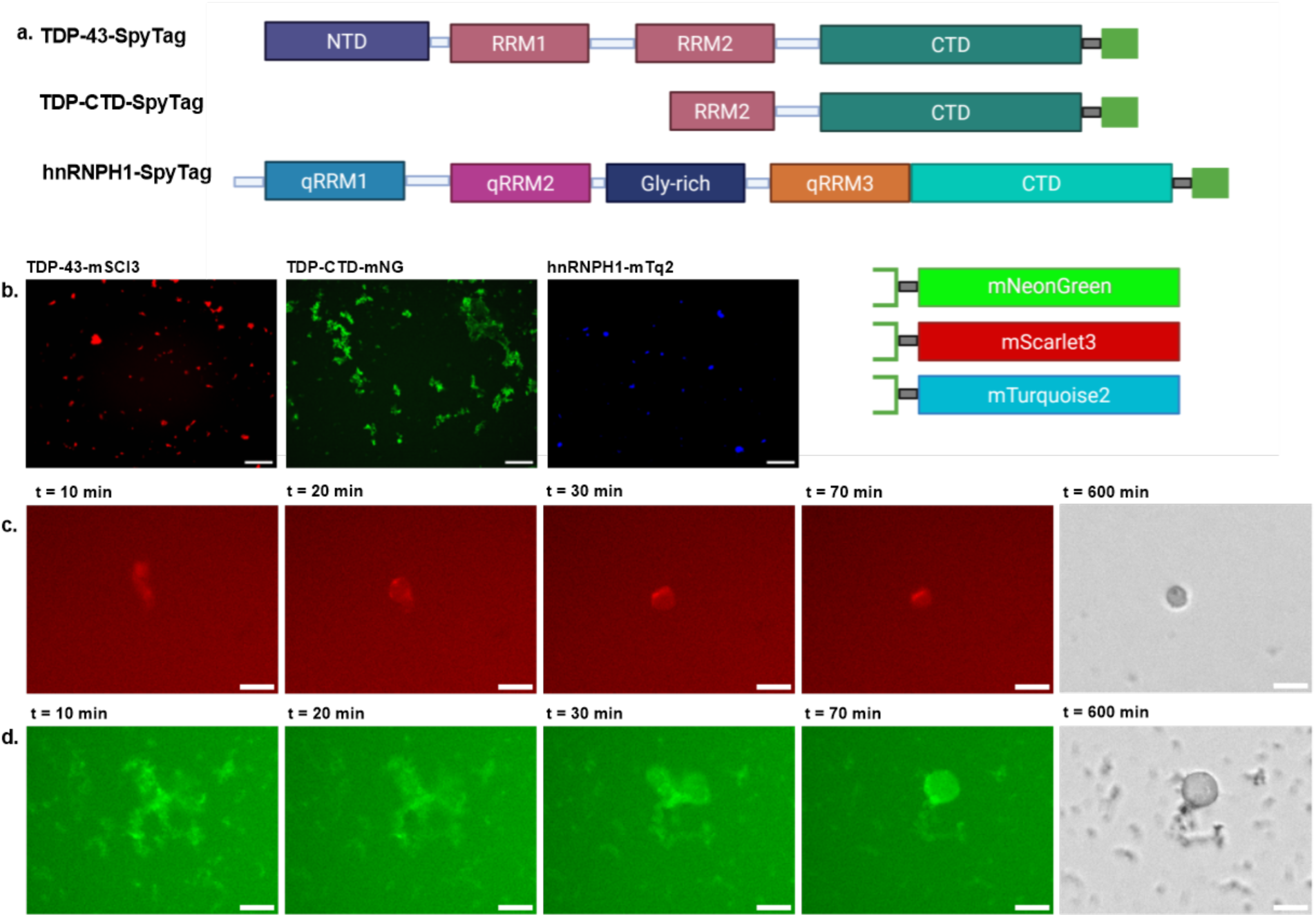
Labeling phase-transition proteins with switchable fluorescent tags. **a** Schematic representation of protein domains fused with SpyTag/SpyCatcher modules. **b** Solid aggregates formed by TDP-43–mScarlet3 (left), TDP-CTD–mNeonGreen (middle), and hnRNPH1–mTurquoise2 (right) under SDS induction conditions. Scale bar is 40μm. **c, d** Time-lapse images showing TDP-43–mScarlet3 (**c**) and TDP-CTD–mNeonGreen (**d**) transitioning from solid aggregates into droplets; the corresponding bright-field image for the final fluorescence frame is shown on the right. Time notations begin after addition of SDS. Scale bars are 10μm.

The pI of all three SpyCatcher-FPs ranges between 5.0 and 5.5. When fused with TDP-43, TDP-CTD, and hnRNPH1, the resulting fusion proteins have pIs between 5.5 and 6.0, predicting low net charges like the untagged proteins under our aggregation induction conditions. As expected, when the buffer was adjusted to pH 6.0 with 0.5M NaCl, the fused proteins formed solid aggregates, exhibiting strong and uniform fluorescence (Fig. 5b). Each protein can be fused to any FP without apparent differences in behavior when the FP was switched among the three monomeric variants (Supplementary Fig. 6). We concluded that adding an FP to the C-terminus of these proteins does not significantly alter their phase behavior, while it affords a convenient and switchable end labeling system to study these proteins. Older versions of green and red FPs have been previously used in fusions to study TDP-43 both in vivo and in vitro^37,53,60^. Our data is consistent with these findings, indicating that FP tags offer reliable information about native proteins. Most commonly used FPs have a slightly acidic pI, making them unlikely to interfere with pH-dependent effects on net charge or electrostatic interactions of proteins with similar pIs as that of TDP-43. However, caution may be warranted about the tendency of conventional avGFP-derived GFP, YFP, and their variations to form weak dimers^61^, particularly when they are used to study oligomerization and seeding for aggregation.

With fluorescence signals appended to each protein, we performed time-lapse imaging to track the transition of individual aggregates of TDP-43 and TDP-CTD that were fused to mScarlet3 and mNeonGreen, respectively. Spread-out aggregates of each protein began condensing within 10 minutes of SDS addition and self-merged into spheroid structures within 1 hour (Fig. 5C), in agreement with results shown with non-fusion TDP-43 (Fig. 1b).

### Electrostatic-driven phase separation is differential to aggregation seeding

Having observed that TDP-43, its CTD, and hnRNPH1—the latter a distinct protein with comparable features to TDP-43—all exhibited parallel behaviors, we explored whether phase separation is directed by protein-specific features, or if it is driven by a common physical property-based process, such as hydrophobic inclusion. We pairwise-mixed all three proteins, each tagged with a different FP, and examined aggregates for the presence of two different proteins. Interestingly, coaggregates were readily observed between full-length TDP-43 and TDP-CTD (Fig. 6). However, in all view areas obtained from repeated attempts, no coaggregates were detected between hnRNPH1 and either TDP-43 or TDP-CTD. Because TDP-CTD is a segment of TDP-43 and known to be required for phase separation^24,26^, we hypothesized that the cocomplex between TDP-43 and TDP-CTD forms through the shared helical seed within the LLPS core previously identified by biochemical and structural studies^14,28^. The LCD core sequence of hnRNPH1 differs from that of TDP-43’s while it is also known to be involved in phase separation^43^. These results suggest that protein assembly is not driven solely by hydrophobic or other nonspecific physical forces, as the process appears protein-specific. Instead, it is plausible that under conditions controlling electrostatic interactions combined with hydrophobic segregation from the aqueous phase, the phase separation appears mediated, or even initiated, at their LCD cores, similar to the concept of “steric zipper”^62^. These flexible regions may allow transient ‘kiss-and-run’ interactions when no matching partner is found, or bundle to form fibrils and develop amyloids when matching occurs. Recently, non-canonical mechanism of protein recognition, wherein transient interactions surrounding the TDP-43 conserved helical region drive the phase separation of CTD through direct enhancement of site-specific, helix-mediated interactions, was illustrate as an example of how the amino acid sequence can be used as a code for segregation^63^. It was previously reported that the mutations within the core of 279-360 (as defined by Kumar *et al.*^28^) rendered TDP-43 less liquid-like^24^, and led to defects in stress granule formation^64^. The sequence specificity in multiple protein assembly formation revealed by our results could be useful for understanding the effects of these mutations.

**Figure 6.**
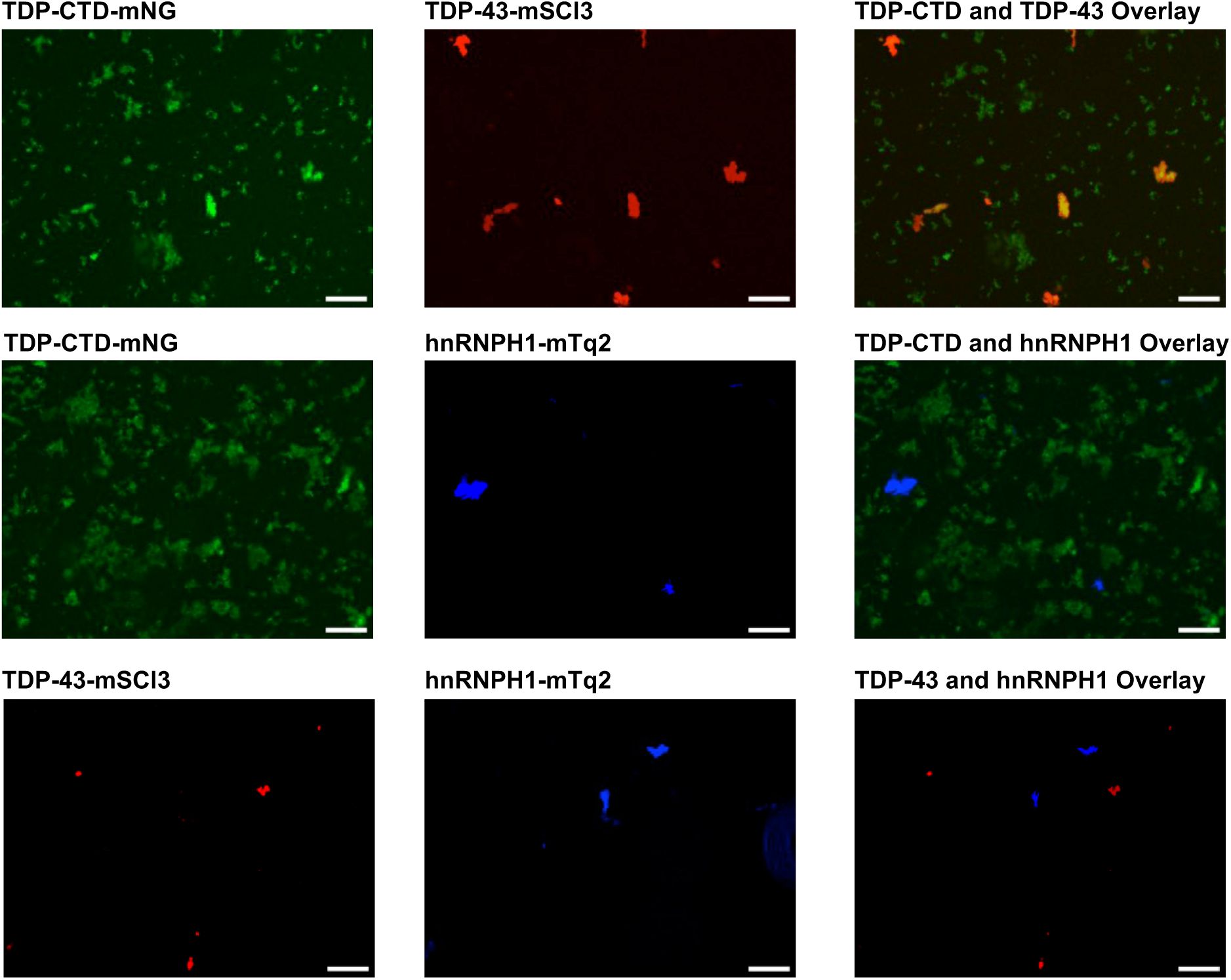
Seeding of phase transition depends on protein compatibility. Pairwise mixing of TDP-CTD–mNeonGreen (mNG), TDP-43–mScarlet3 (mSCI3), and hnRNPH1–mTurquoise2 (mTq2) in solution prior to induction to form aggregates. Coaggregates between TDP-CTD–mNeonGreen and TDP-43–mScarlet3 appear as yellow-orange due to fluorescence overlap. Scale bar is 20μm.

### Toward modulation of phase transition

Based on the experimental data presented in this study, we propose a unifying perspective about phase behavior of proteins such as TDP-43. Electrostatics related to charge, whether global or localized, arising from natural amino acid composition and post-translational modifications is a major determinant. In vitro attunement to pH, salt screen, zwitterionic glycine (and other amino acids) or charge decoration by ionic detergent SDS at very low concentration, can modulate how proteins like TDP-43 transition among various macroscopic phases. These manipulations can induce phase transitions by shifting some microscopic substructures such as dimers, oligomers, nonamyloid fibrils and amyloids to reach the lowest-energy steady state. These substructures all retain a degree of fluidity and reversibility, allowing the system to respond dynamically to environmental changes.

Protein assemblies can be visualized at the micron scale using optical microscopy, as shown in this study, or at the nanometer scale with electron microscopy and other high-resolution techniques like NMR, revealing details such as filament-containing helical amyloid core^28^. Large assemblies likely originate from smaller subunits that act as building blocks, beginning as oligomers and progressing through stepwise assembly. The process typically takes time, occurring over hours or days^31^, though notably faster in our system. These substructures may share physical and chemical characteristics with the larger assemblies they form, either through fractal growth into amorphous solids or by coexisting in a milieu that phase-separates into liquid-like droplets. In our system, strong ThT and CR staining was observed in both solids and droplets, during and after solid-to-droplet transitions with no delays, suggesting that different forms might share related substructures. These staining events may originate from a seed region containing an amphipathic helix shared among aggregating molecules^29^, which undergo conformational transitions toward increased β-sheet stacking during maturation, under stress, or with mutations, as previously proposed^30,49,65^. Such seeding could allow the initial helical core and adjacent sequences to act as a sorting mechanism, determining interaction partners and conditions. It is important to note that RNA-binding proteins can also interact between each other as soluble molecules via LCDs, such as glycine-rich regions, to perform biological functions without undergoing phase transitions, as shown in our earlier in vitro protein-protein interaction studies^66,67^.

TDP-43 is known to co-segregate under physiological conditions with numerous other proteins, including hnRNPA1, hnRNPA2, and FUS^3,68,69^, through domain-specific physical contact, via RNA binding, or perhaps coaggregation based on matching helix or β-sheet stacking at seed region. The phase transitions observed here with TDP-43 and hnRNPH1 may extend beyond RNA-binding proteins involved in splicing regulation. A notable early example is Sear’s 2007 prediction that the Dvl protein would undergo LLPS^70^, an all-or-none switch that has since been shown to play a critical role in the Wnt signaling pathway^71^.

The discovery that solid aggregates can be driven into a more liquid-like state may open new avenues for identifying therapeutic intervention points by preserving normal, reversible LLPS that does not progress to irreversible fibril amyloid formation^72^. Potential strategies include using short RNAs to provide specificity by targeting RNA-binding proteins, and amino acids or oligopeptides to achieve charge-modifying effects.

## Materials and Methods

### Protein production

Plasmids encoding TDP-43, TDP-CTD, hnRNPH1, and SUMO–TDP-43 were cloned into the pNCS vector^57^ and transformed into NEB SHuffle E. coli competent cells. Selected colonies were inoculated into culture tubes containing 5mL 0.1 mg/mL carbenicillin 2xYT broth and incubated at 30°C overnight while shaking, the next day transferred to 500mL 2xYT broth and again incubated overnight while shaking. For expression, small cultures of transformed bacteria were inoculated in 500 mL 2×YT broth supplemented with 0.1 mg/mL carbenicillin at 30°C while shaking overnight. Cells were harvested by centrifugation at 8,000 × g, and the pellets were resuspended in lysis buffer containing 20 mM Tris-HCl (pH 8.0), 6 M urea, 500 mM NaCl, and 10 mM imidazole.

Cells were lysed by sonication on ice (30% amplitude, 5 min total, 10 s on/10 s off) and centrifuged at 8,000 × g to remove debris. The supernatant was incubated at 4 °C for 1 h to overnight with nickel beads to capture His-tagged proteins. Bead–protein complexes were transferred to purification columns and washed 3–4 times with wash buffer (20 mM Tris-HCl, pH 8.0; 500 mM NaCl; 10 mM imidazole; 6 M urea). Proteins were eluted in a single step using elution buffer (20 mM Tris-HCl, pH 8.0; 500 mM NaCl; 200 mM imidazole; 4.5 M urea) and filtered through 0.22 µm PES syringe filters.

Protein purity was assessed by SDS-PAGE, and concentrations were measured using NanoDrop. Eluted proteins were standardized to 5 µM, or about 0.2 mg/ml, after removal of urea in gradual 5-6 step dialysis using Spectra/Por dialysis tubing (MW cutoff 7,000) against storage buffer of 20 mM Tris-HCl, pH 8.0 100 mM NaCl with urea concentration decreasing ∼50% per step until all urea was removed. Soluble proteins were collected and stored at −20 °C.

### Formation of solid aggregates

Phase separation reactions for TDP-43, TDP-CTD, and hnRNPH1 were initiated from soluble protein fractions by adjusting the storage buffer from pH 8 to pH 6 using small-volume additions of 1 M Tris-HCl pH buffer. Additional modifications, such as changes in salt concentration, were made by adding small volumes of high-molar salt solutions. The total reaction volume was approximately 40 µL. All added reagents to modify experimental conditions resulted in <10% increase to the total reaction volume. Aggregates from all proteins were monitored at designated time points, with complete formation typically occurring within 5 hours.

### Formation of liquid-like condensates

Liquid-like condensates were generated from either soluble proteins or pre-formed solid aggregates by adding 1% SDS directly to the reaction mixture. The transition typically completed within 8 hours. The final SDS concentration was 0.03%, consistent across all stated methods.

In experiments where glycine was added to probe the effects of addition, the final concentration of glycine was 50 mM for experiments shown in Fig. 2c, and 5mM in Fig. 4a. Concentrated glycine was diluted directly to the experimental conditions.

To quantify droplet shape characteristics, three geometric descriptors were calculated from binary images: circularity, roundness, and solidity.

Circularity was calculated using the formula:

Circularity = (4 × π × Area) / (Perimeter²)

This metric ranges from 0 to 1, where a value of 1 indicates a perfect circle. Lower values suggest more elongated or irregular shapes.

Roundness was defined as:

Roundness = (4 × Area) / (π × (Major Axis Length)²)

Roundness also ranges from 0 to 1, with higher values indicating shapes closer to a perfect circle, but it is more sensitive to elongation than circularity.

Solidity was computed as:

Solidity = Area / Convex Hull Area

Solidity measures the extent to which a shape is convex. A value of 1 indicates a completely convex shape, while lower values suggest the presence of concavities or irregular boundaries.

All measurements were performed using image analysis software capable of extracting shape descriptors from segmented droplet images.

### Imaging

Microscopy imaging included bright field, phase contrast, widefield fluorescence, and confocal fluorescence modalities. The primary instrument was the Keyence BZ-X800 fluorescence microscope for both light and fluorescence imaging. Additional bright-field and widefield fluorescence images were captured using the EVOS FL Fluorescence Digital Microscope, while confocal fluorescence imaging was performed with the BioTek Cytation C10 Confocal Imaging Reader. Images taken by Cytation C10 were at 40x magnification. All other images were taken at 20x magnification.

### Image analysis and quantifications

Image contrast and brightness were adjusted uniformly for clarity using ImageJ. Quantitative measurements were also performed using ImageJ’s built-in analysis functions.

For quantifications of fluorescence images, ImageJ’s color threshold function was used to delineate positive signals from background signal. The analysis function provided a count of detected structure, the area covered by structures of the total visible area Area%, as well as fluorescence intensity IntDen.

IntDen or Integrated Density is the product of mean gray value (a stand-in for intensity) and area, representing the amount of dye bound and fluorescing. The units are (relative fluorescence units) * (µm^2^). Corrected Total View Fluorescence is used to represent the intensity of fluorescence in our measurements, which takes the IntDen value from each measured object and corrects for background fluorescence. The equation is as follows:

CTFV = IntDen – (Area of Droplet * Mean fluorescence of background)

The mean fluorescence of background was measured by averaging 3 separate readings of the background for each time point and view of the well.

### Dye staining

Protein assemblies were stained by incubating at room temperature for 2–3 hours with Thioflavin T (ThT) or Congo Red (CR) from Sigma. Final concentrations in wells were 40 µM ThT or 0.5% CR. For time course experiments shown in Fig. 4, ThT was added to the solution before the phase separation was induced.

### SpyTag/SpyCatcher Conjugation

Protein constructs were conjugated by incubating together at room temperature for 1 hour. FP–SpyCatcher was added in slight excess (10%) over TDP/CTD/hnRNPH1–SpyTag to achieve a 1.1:1 molar ratio. For quenching, excess SpyTag-tagged nanobodies (at least 4× the molarity of SpyCatcher–FP) were added to bind any remaining free SpyCatcher–FP, followed by an additional 1-hour incubation at room temperature.

## Acknowledgment

We thank Alex Ward for some of the initial DNA constructions and initial attempts at protein production, Andrew Chammas and Nathan Shaner for help with designing of SpyTag and SpyCatcher constructs, Aryan Mudliar for lab assistance and discussions.

## Author Contributions

JW conceptually initiated the project; TRW designed and carried out the experiments; TRW and JW interpreted data and drafted the manuscript.

## Data Availability

Additional, related data of the present report is available from the corresponding author upon reasonable request.

## Supplementary Material

Supplementary material.doc: Supplementary figures, materials and methods

