## Supplemental Material for "Phase Behavior of TDP-43 and hnRNPH1: From Soluble and Aggregated States to Liquid-Like Droplets" (from another version of this preprint: version 2, 2025-11-14)

### Supplementary Figure 1

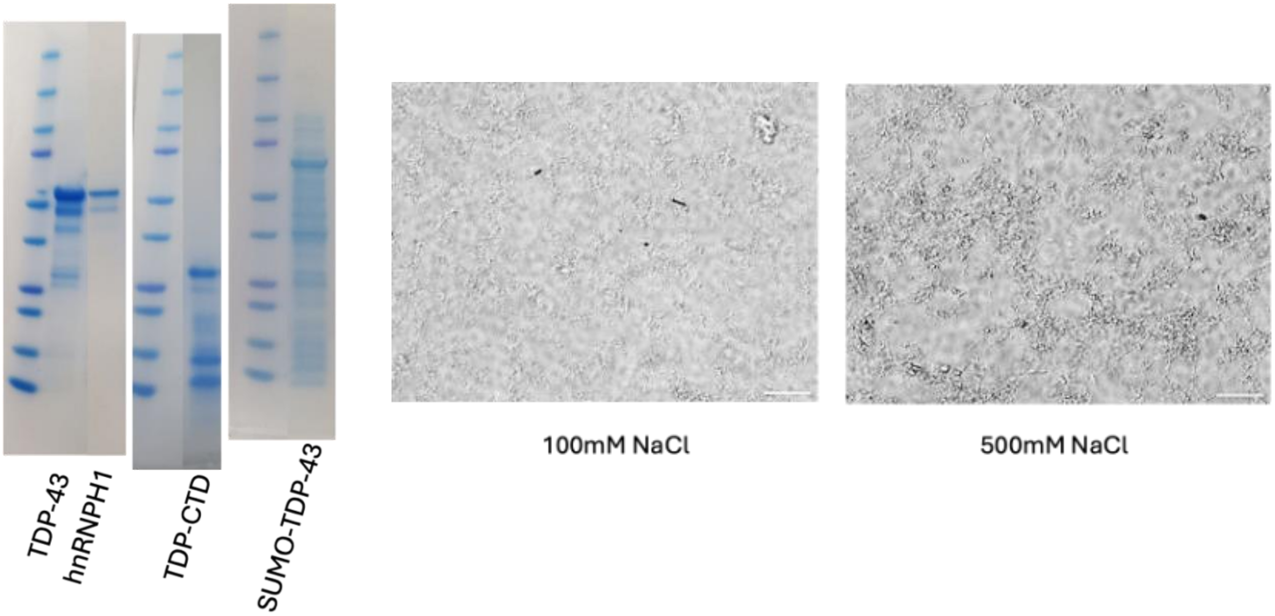


**Recombinant proteins and conditions for inducing phase separation**

**a** Full-length TDP-43 forms aggregates under both low (100 mM) and high (500 mM) NaCl conditions (right), with high salt favoring phase separation. Scale bars are 40µm. SDS-PAGE gel showing purified protein bands; minor bands below the expected TDP-43 size were commonly observed, as previously shown [Staderini, 2022 #242] (right). **b** Aggregate formation at different pH values with ThT added (as in Fig. 3 of the main text) for enhanced visualization. pH 6 promotes more extensive aggregation. Scale bar is 20µm.

### Supplementary Figure 2

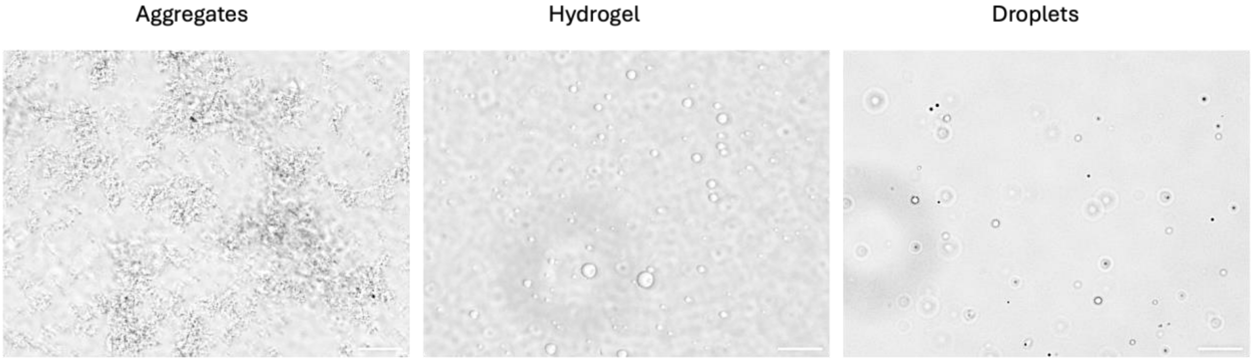

**SUMO tag does not significantly alter TDP-43 phase separation**

Phase separation of SUMO-tagged TDP-43, which has a similar pI to untagged TDP-43, forming solid aggregates (left) at pH 6 and 500 mM NaCl; droplets formed directly with 0.03% SDS (middle); and droplets induced from aggregates upon SDS addition (right). All behaviors are comparable to untagged TDP-43 as shown in Figure 1. Scale bars are 40μm.

### Supplementary Figure 3

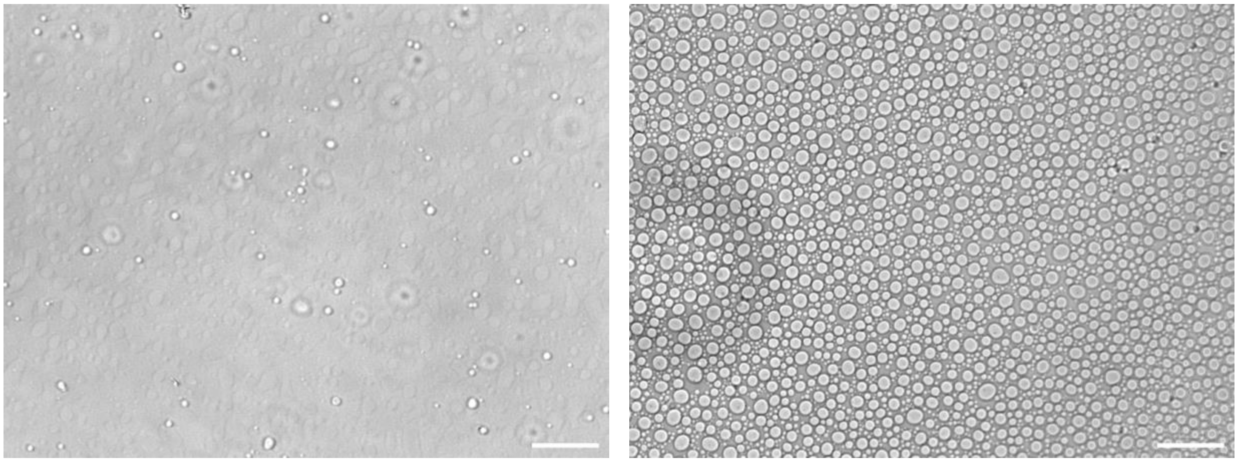

**Crowding agent enhances solution-to-droplet LLPS**

Addition of PEG8000 (right) markedly increases droplet formation induced by SDS compared to the control condition without PEG (left). Scale bars are 40μm.

### Supplementary Figure 4

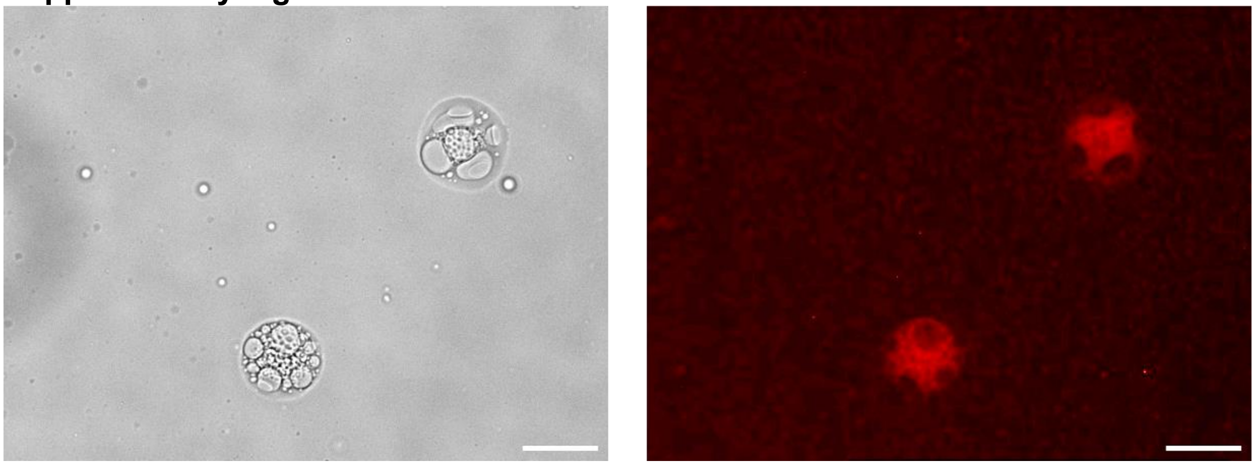

**mRNA remains associated with TDP-43 during phase separation**

Cy5-labeled mRNA of fluorescent protein mWasabi as irrelevant mRNA to a final concentration of 15ng/µL was added to soluble TDP-43, and multi-emulsion droplets were induced as described in the main text. Shown are bright-field (left) and fluorescence (right) images of a representative multiphase droplet, where mRNA signal overlaps with the dense phase. This indicates that protein-bound mRNA can enter multiphase droplets together with TDP-43. Scale bars are 10µm.

### Supplementary Figure 5

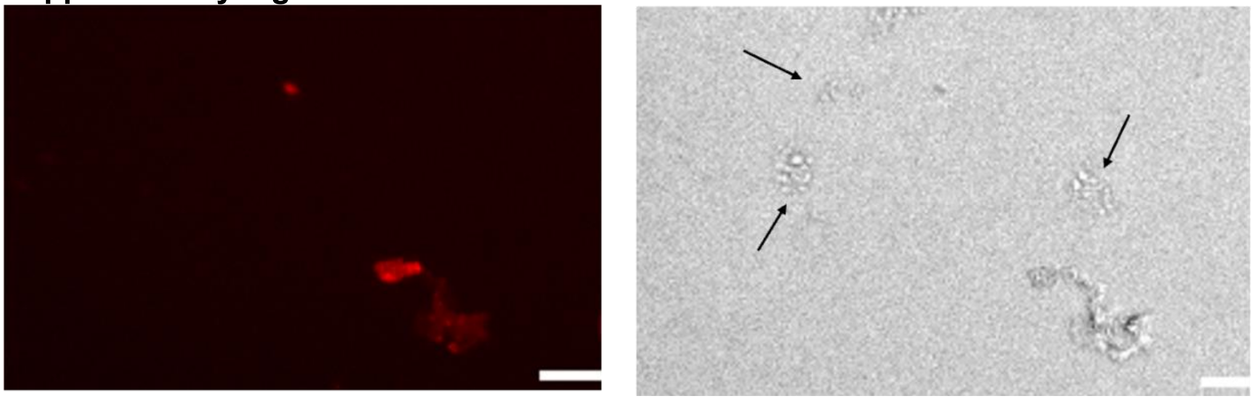

**SpyCatcher-FP can be quenched**

To ensure SpyCatcher does not carry over into coaggregation experiments, an unrelated Spy-tagged nanobody (14 kDa, non-aggregating) was used as a quencher after the labeling step. Shown are aggregates that remained labeled (solid arrow) and aggregates left unlabeled after SpyCatcher was quenched (open arrow).

### Supplementary Figure 6

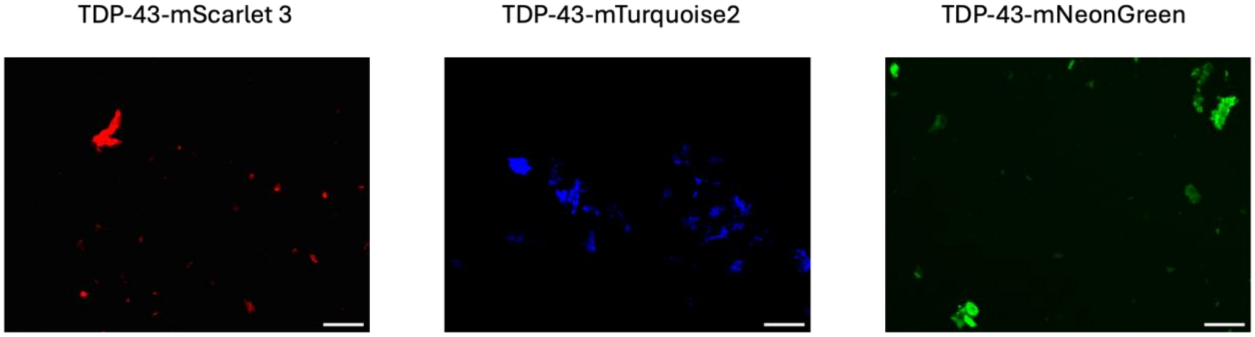

**SpyTag–TDP-43 can be labeled with multiple fluorescent colors using different SpyCatcher–FPs**

Shown are TDP-43 aggregates labeled in three distinct colors after fusing TDP-43–SpyTag with different SpyCatcher–FP constructs, as indicated in the figure. Scale bars are 20μm.

### Supplement Materials and Methods

#### SDS-PAGE

Protein samples were run on 20% MINI-PROTEAN TGX gels. Loading dye consisted of a 1:10 dilution of 2-mercaptoethanol to 5x non-reducing lane marker sample buffer. 2μL loading dye was added to 10μL of protein before being heated to 98°C for 5 min for denaturation. Electrodes were set to separate the samples at 220V for 20-23 minutes. Following removal from the gel rig, gels were washed in distilled water and shaken for 5 minutes before staining in Bio-Rad Bio-Safe™ Coomassie Stain for a minimum of 1 hour. After staining, the dye was safely disposed of and destained in distilled water overnight prior to imaging.

#### PEG addition

Droplets of Sumo-TDP-43 were generated from soluble solution as described above. The effects of PEG were studied by addition of PEG-8000 to a final 0.7% within the reactions. Pictures were taken after the reaction appeared to reach stable equilibrium within 1 hour.

#### mRNA addition

mRNA of mWasabi fluorescent protein designed to stain in Cy5 and adjacent channels was added to a final concentration of 15ng/μL in the initial solution of Sumo-TDP-43 before appearance of insoluble structures. 6-7 hours following induction with SDS, pictures were of stained droplets in brightfield and RFP filters.
